# Daily administration of low-dose daunorubicin or doxorubicin inhibits hypoxia-inducible factor 1 and tumor vascularization

**DOI:** 10.1101/2022.06.15.492526

**Authors:** Yongkang Yang, David Z. Qian, Sergio Rey, Jun O. Liu, Gregg L. Semenza

## Abstract

Using a hypoxia-inducible factor 1 (HIF-1)-dependent luciferase reporter in Hep3B human hepatocellular carcinoma cells, we screened over 3,000 drugs that have been used in clinical trials and identified multiple anthracyclines as inhibitors of HIF-1 activity. Anthracyclines interfered with the ability of HIF-1 to bind to DNA. Daily injection of tumor-bearing mice with anthracyclines at low dose inhibited expression of the luciferase reporter and HIF-1 target genes that encode vascular endothelial growth factor A (VEGFA; ligand of VEGFR2), stromal-derived factor 1 (SDF-1; ligand of CXCR4), and stem cell factor (SCF; ligand of CD117) in tumor tissue. Increased numbers of circulating CXCR4^+^/Sca1^+^, VEGFR2^+^/CD34^+^, and VEGFR2^+^/CD117^+^ cells were demonstrated in immunodeficient mice bearing prostate cancer xenografts but not in tumor-bearing mice treated with anthracyclines, which also significantly inhibited angiogenesis in tumor tissue. Our findings indicate that HIF-1 inhibition underlies the anti-angiogenic effect associated with daunorubicin or doxorubicin metronomic therapy and suggest that these drugs may be particularly effective in patients with high levels of HIF-1α in their diagnostic tumor biopsy.

## Introduction

Many advanced human cancers are characterized by regions in which the partial pressure of O_2_ (pO_2_) is significantly decreased compared to normal tissue and the presence of severe hypoxia (pO_2_ < 10 mm Hg by direct Eppendorf microelectrode measurement) is associated with increased risk of tumor invasion, metastasis, and patient mortality (1, 2). The major mechanism by which decreased intratumoral pO_2_ promotes cancer progression is through increased expression and activity of HIF-1, which is a transcriptional activator that regulates the expression of hundreds of genes that encode proteins involved in cancer stem cell specification, energy metabolism, immune evasion, invasion and metastasis, treatment resistance, and tumor vascularization (3). HIF-1 transactivates the *VEGFA* gene, which encodes vascular endothelial growth factor A (4). Genetic or pharmacological HIF-1 gain-of-function or loss-of-function in human cancer cells has been shown to increase or decrease, respectively, VEGFA expression and tumor angiogenesis (1, 5-10). HIF-1 also activates transcription of genes encoding glucose transporters (GLUT1 and GLUT3) and hexokinases (HK1 and HK2), which mediate the high level of glucose uptake and phosphorylation that is observed in cancer cells, and lactate dehydrogenase A and pyruvate dehydrogenase kinase 1 (PDK1), which shunt pyruvate away from the mitochondria and increase lactate production (1, 7, 11, 12). Through the regulation of these and other genes controlling angiogenesis and glucose metabolism, HIF-1 modulates both O_2_ delivery and O_2_ consumption, respectively.

HIF-1 is a heterodimer consisting of HIF-1α and HIF-1β subunits (13). HIF-1α levels increase exponentially as O_2_ availability declines (14). In normoxic cells, the human HIF-1α protein is subjected to hydroxylation on proline residue 402 and/or 564, which is required for binding of the von Hippel–Lindau tumor suppressor protein, which is the recognition subunit of a ubiquitin ligase complex that marks HIF-1α for proteasomal degradation (15, 16). In normoxic cells, asparagine residue 803 of human HIF-1α is also hydroxylated, which blocks the binding of coactivator proteins p300 and CBP to the transactivation domain (TAD) of HIF-1α (17). The hydroxylases use O_2_ as a substrate and their activity declines under hypoxic conditions, leading to increased HIF-1α stabilization and transactivation. HIF-1 binds to hypoxia response elements (HREs), which contain the consensus sequence 5’-(A/G)CGTG-3’ (12) and can be located within the body of a HIF-1 target gene or in its flanking sequences.

Increased HIF-1α protein expression in cancer biopsy sections is associated with patient mortality in a wide variety of cancer types (18). Inhibitors of HIF-1 may be useful for cancer therapy and compounds that inhibit HIF-1α synthesis, protein stability, subunit heterodimerization, DNA binding activity, or TAD function have been reported (19). In this study, we found that anthracyclines, which constitute one of the most well-known classes of chemotherapy drugs, interfere with HIF-1 binding to DNA and that inhibition of HIF-1 activity provides a mechanism of action for the anti-angiogenic effects of these drugs.

## Results

### Anthracyclines Inhibit HIF-1 Transcriptional Activity

To screen for low molecular weight compounds that inhibit HIF-1 activity, we developed a reporter assay utilizing a human hepatocellular carcinoma line Hep3B subclone (designated Hep3B-c1) that is stably transfected with plasmid p2.1, in which firefly luciferase (FLuc) coding sequences are under the control of an HRE from the human enolase gene upstream of a basal SV40 promoter (12), and plasmid pSV- Renilla, in which Renilla luciferase (RLuc) coding sequences are under the control of the SV40 promoter only (10). The FLuc:RLuc ratio was more than three-fold higher when Hep3B-c1 cells were exposed to hypoxic conditions (1% O_2_) as compared to non-hypoxic conditions (20% O_2_) (Figure 1A, upper panels, and Figure S1A). We screened a collection of 3,120 drugs that have been approved by the U.S. FDA or have been evaluated in phase I clinical trials (20). The most potent inhibitor of HIF-1-dependent reporter gene expression identified by this assay was daunorubicin (DNR), which inhibited hypoxia-induced FLuc:RLuc activity by 99% at a concentration of 10 μM. Two other anthracyclines, epirubicin (EPI) and idarubicin (IDA), were also identified in the Hep3B reporter assay. DNR, EPI, IDA, and doxorubicin (DXR) inhibited hypoxia-induced FLuc:RLuc activity at concentrations as low as 200 nM in a dose-dependent manner (Figure 1A, upper panels, and Figure S1A). The anthracyclines also inhibited the expression of endogenous HIF-1 target genes: VEGF and GLUT1 mRNA levels were decreased by treatment of hypoxic cells with anthracyclines in a dose-dependent manner (Figure 1A, middle and lower panels, and Figure S1B). Increased FLuc:RLuc activity that was mediated by cotransfection of expression vector encoding HIF-1α or HIF-2α was also inhibited by anthracycline treatment (Figure 1B). Taken together, the results presented in Figure 1 and Figure S1 demonstrate that anthracyclines inhibit HIF-1-dependent transcription in hypoxic cancer cells.

**Figure 1.**
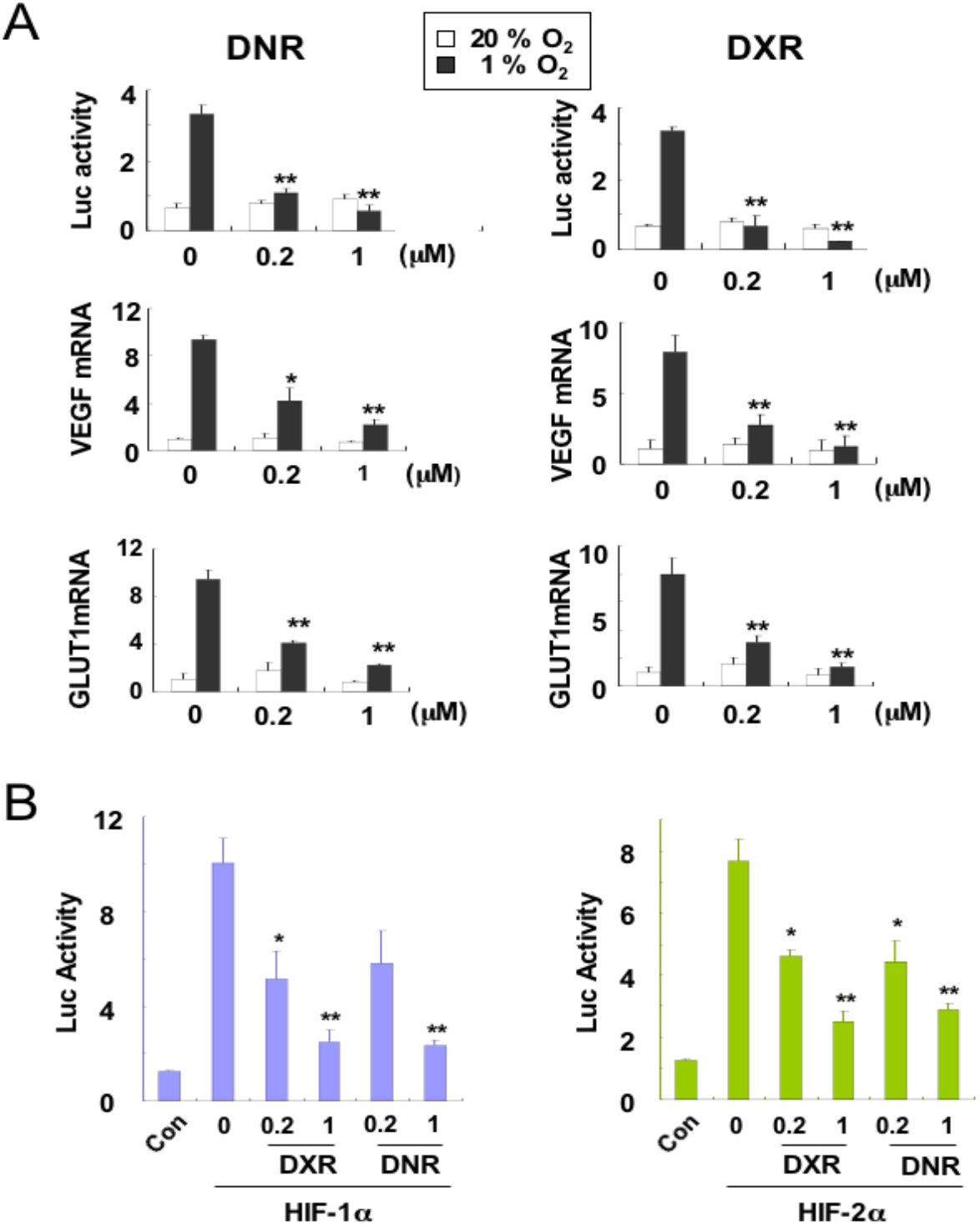
Anthracyclines inhibit gene expression mediated by HIF-1 and HIF-2. (**A**) Treatment of cells with daunorubicin (DNR) or doxorubicin (DXR) inhibits hypoxia-induced gene expression. *(Top panel)* HIF-1-dependent luciferase activity (FLuc:RLuc ratio) was analyzed in Hep3B-c1 cells treated with the indicated concentration of DNR *(left panel)* or DXR *(right panel)* and exposed to 20% O_2_ (white bars) or 1% O_2_ (black bars) for 24 hours. *(Middle and bottom panels*) HEK-293 cells were exposed to 20% O_2_ (white bars) or 1% O_2_ (black bars) for 24 hours in the presence of the indicated concentration of DNR or DXR and total RNA was isolated for determination of VEGFA mRNA *(middle panel)* and GLUT1 mRNA *(lower panel)* levels by reverse transcription and real-time quantitative PCR (RT-qPCR). The mRNA levels were normalized to the levels of 18S rRNA in each sample, and expressed relative to the levels in vehicle-treated cells exposed to 20% O_2_. Mean ± SEM are plotted (n = 4). *, *P* < 0.05; **, *P* < 0.01. (**B**) Treatment of HEK-293 cells with DNR or DXR inhibits luciferase activity (FLuc:RLuc ratio) mediated by forced expression of HIF-1α *(Left, blue bars)* or HIF-2α *(Right, green bars).* Mean ± SEM are plotted (n = 3). *, *P* < 0.05; **, *P* < 0.01.

### Anthracyclines Do Not Affect HIF-1α Protein Levels

To establish the molecular mechanism by which DNR, DXR, EPI, and IDA inhibit HIF-1-dependent transcription, we first analyzed HIF- 1a expression by immunoblot (IB) assay. Stabilization of HIF-1α protein is a critical mechanism underlying increased HIF activity in response to hypoxia and several chemical compounds inhibit HIF-1 activity by causing HIF-1α degradation (19). Cells were exposed to 20% O_2_ or to 1% O_2_ in the presence of vehicle or anthracycline for 20 hours. HIF-1α protein was barely detectable in cells exposed to 20% O_2_ and was greatly increased in cells exposed to 1% O_2_ (Figure S2A). The hypoxia-induced expression of HIF-1α was not inhibited by anthracycline treatment.

### Anthracyclines Do Not Affect HIF-1α TAD Function

To determine whether anthracyclines impair the function of the HIF-1α TAD, which is located in the carboxyl terminal half of the 826- amino-acid HIF-1α protein, cells were co-transfected with: pSV-Renilla; an FLuc reporter gene containing five GAL4-binding sites; and an expression vector encoding the yeast GAL4 DNA-binding domain (DBD), either alone or fused to HIF-1α sequences (21). In cells expressing GAL- O (GAL4-DBD alone), FLuc:RLuc activity was low and was not induced by hypoxia (Figure S2B). By contrast, expression of GAL-A (GAL4-DBD fused to HIF-1α residues 531–826) or GAL-C (GAL4-DBD fused to HIF-1α residues 653–826) resulted in higher FLuc:RLuc activity at 20% O_2_ and a greater than 6-fold increase in FLuc:RLuc activity in response to hypoxia (Figure S2B). FLuc:RLuc activity was not affected by anthracycline treatment. GAL-H (GAL4-DBD fused to HIF-1α residues 786–826), which lacks the binding site for the asparagine hydroxylase FIH-1 and is constitutively active (17, 21, 22), was also not inhibited by treatment with DNR, DXR, IDA, or EPI (Figure S2B). Based on these results, we conclude that anthracyclines do not directly inhibit HIF-1α TAD function.

### Anthracyclines Do Not Affect HIF Subunit Dimerization

We next investigated whether anthracyclines inhibit the heterodimerization of HIF-1α and HIF-1β, which is required for DNA binding and target gene activation. We expressed a glutathione-*S*-transferase (GST)-HIF-1β fusion protein in bacteria and purified the protein by capture on glutathione-agarose beads, which were then incubated with lysates from HEK-293 cells transfected with a vector encoding FLAG epitope-tagged HIF-1α. FLAG-HIF-1a was pulled down by GST-HIF-1β but not by GST alone, and this protein-protein interaction was not affected by anthracycline treatment (Figure S2C). Based on these results, we conclude that anthracyclines do not block HIF-1 activity by inhibiting the heterodimerization of HIF-1α and HIF-1β.

### Anthracyclines Inhibit Binding of HIF-1 to DNA

Another potential molecular mechanism by which anthracyclines block HIF-1-dependent transcription is by inhibiting the binding of HIF-1 to DNA. To test this hypothesis, we performed chromatin immunoprecipitation (ChIP) assays. Chromatin-bound HIF-1α was immunoprecipitated from HEK-293 cells, which had been exposed to 1% O_2_ or 20% O_2_ for 20 hours in the presence of vehicle or anthracycline. HIF-1 binding sites located in the *VEGFA* and *PDK1* genes were specifically amplified by PCR using chromatin immunoprecipitated from vehicle-treated hypoxic cells, demonstrating the hypoxia-induced binding of HIF-1 to the HRE in these genes (Figure 2). When the cells were treated with anthracycline, binding of HIF-1 to these DNA sequences was inhibited. It is known that anthracyclines bind to DNA, with optimal target sequences of 5’-(A/T)CG-3’ and 5’-(A/T)GC-3’ (23, 24), which overlap with the HIF-1 binding site sequence 5’-(A/G)CGTG-3’ (12). Exposure of cells to DNR or DXR also inhibited binding of HIF-2α to the *VEGFA* and *PDK1* genes in hypoxic cells (Figure S3).

**Figure 2.**
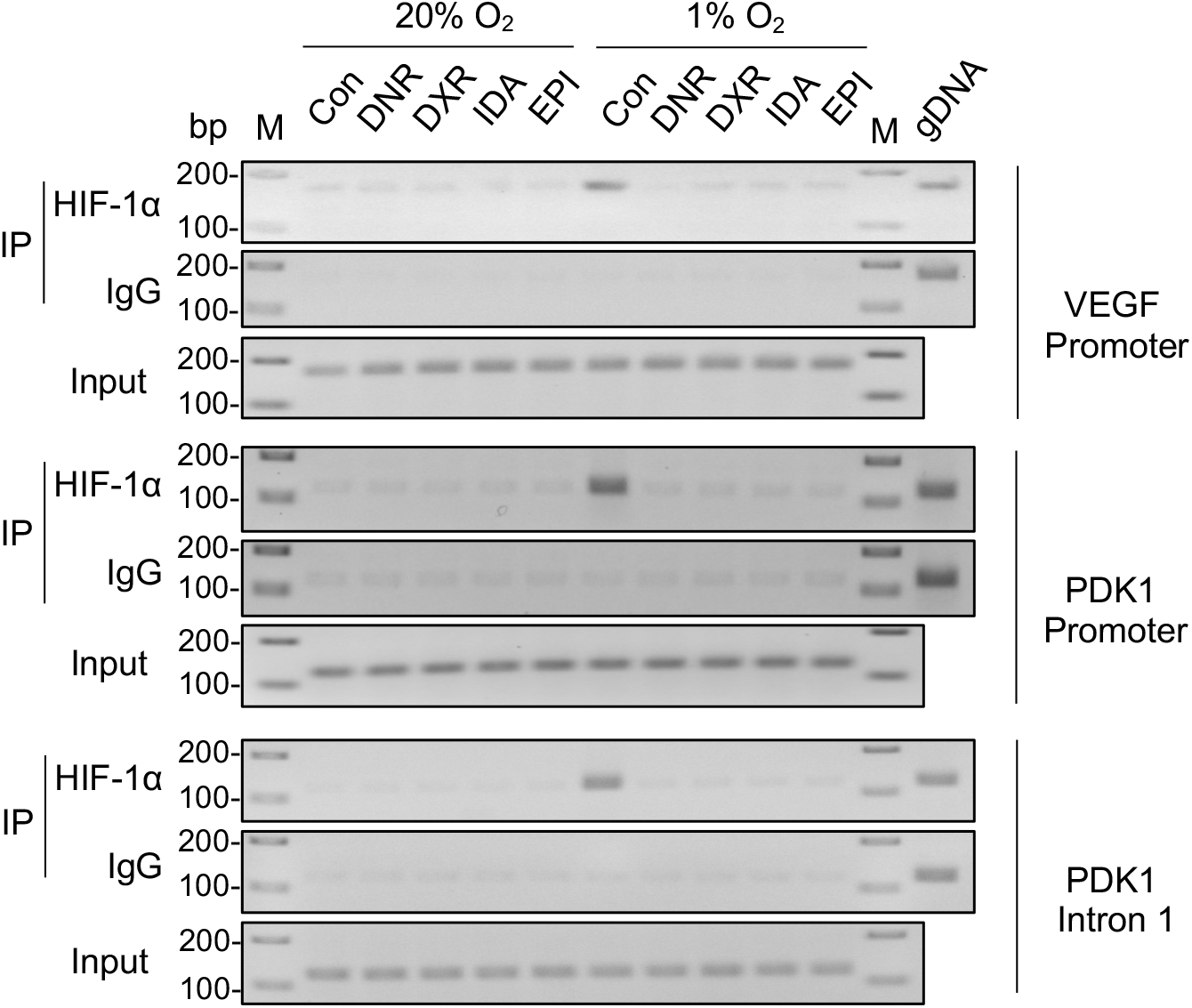
Anthracyclines inhibit hypoxia-induced binding of HIF-1 to target genes. HEK-293 cells were exposed to 20% O_2_ or 1% O_2_ for 20 hours in the presence of vehicle control (Con) or the indicated anthracycline at a concentration of 1 μM. Cell lysates were prepared and aliquots were immunoprecipitated with an antibody against HIF-1α or with rabbit IgG. PCR was performed using immunoprecipitated chromatin as the template for amplification of *VEGFA* promoter, *PDK1* promoter, and *PDK1* intron 1 sequences that each encompass a HIF-1 binding site. An aliquot of chromatin was reserved before immunoprecipitation for PCR (Input). Purified genomic DNA was also subject to PCR (gDNA). M, DNA ladder.

### Anthracyclines Block HIF-1 Activity In Vivo

Anthracyclines are used in the treatment of leukemias, lymphomas, sarcomas, and carcinomas (25). To investigate whether inhibition of HIF- 1 transcriptional activity by anthracyclines contributes to their therapeutic effect, we analyzed the effect of anthracycline treatment on GLUT1 and VEGFA mRNA expression in Hep3B-c1 tumor tissue. We first demonstrated that these anthracyclines did not alter HIF-1α protein levels in cultured Hep3B-c1 cells (Figure S4A), whereas HIF-1 target gene expression was inhibited by anthracycline treatment of hypoxic Hep3B-c1 cells (Figure S4B), which was similar to the effects of these drugs in HEK-293 cells (Figure 1).

Next, five million Hep3B-c1 cells were injected subcutaneously into immunodeficient mice. When the tumors reached 200 mm^3^, the mice were randomly assigned to treatment with DNR, DXR, or vehicle. Prior to treatment (day 27), mean tumor volumes were similar in the three groups (Figure 3A). Analysis of FLuc activity in the tumor by Xenogen imaging revealed that bioluminescence was also similar in the three groups (Figure 3B, upper panel). Anthracycline or vehicle was injected intravenously on four consecutive days at a dose of 1 mg/kg, which is five times lower than the anthracycline dose that is conventionally used to inhibit tumor xenograft growth (26). HIF-1-dependent FLuc activity was imaged 4 hours after the third injection. The bioluminescence in tumors before (day 27) and after (day 30) treatment with vehicle was similar. By contrast, FLuc activity was significantly decreased after anthracycline administration (Figure 3B). The tumor volume on day 30 did not differ significantly between treatment groups (Figure 3A), indicating that inhibition of HIF-1 reporter gene expression by anthracycline administration preceded the inhibitory effect on tumor growth on day 31.

**Figure 3.**
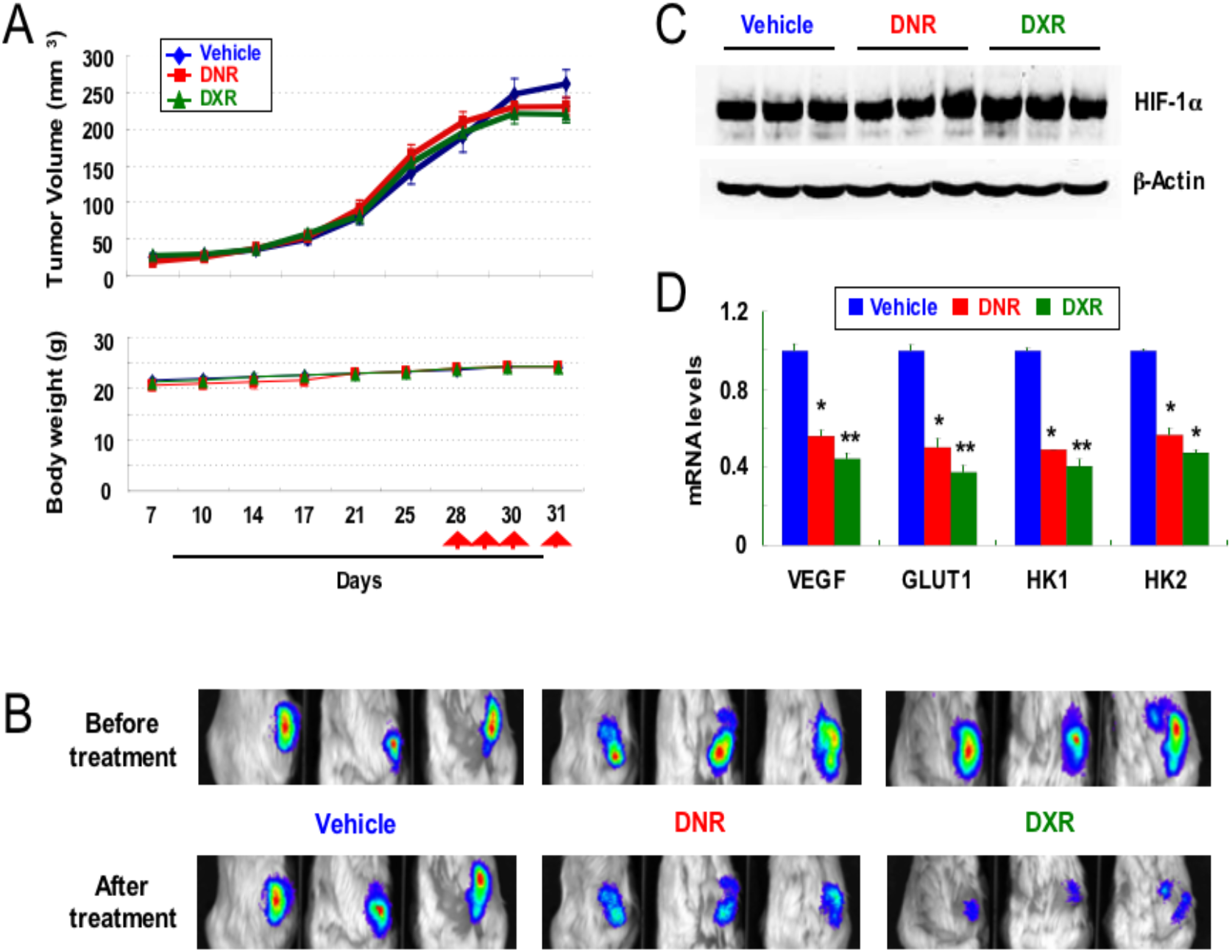
Anthracycline treatment inhibits HIF activity in tumor xenografts. (**A**) Mice were injected subcutaneously with Hep3B-c1 cells and when tumors reached a volume of 200-mm^3^ the mice were administered vehicle (blue), DNR (1 mg/kg/day, red), or DXR (1 mg/kg/day, green) intravenously on days 28-31 (red arrows). Tumor volume and body weight were monitored twice weekly. (**B**) HIF-1-dependent FLuc activity was determined by Xenogen imaging before treatment on day 27 *(top panels)* and 4 hours after last dose on day 31 *(bottom panels).* (**C** and **D**) Tumor xenografts were harvested 4 hours after treatment on day 31 for analysis of HIF-1α protein levels by immunoblot (IB) assay (**C**) and levels of the indicated mRNAs by RT-qPCR (**D**). Data are presented as mean ± SEM (n = 3). *, *P* < 0.05; **, *P* < 0.01.

IB assays of tumors harvested 4 hours after administration of the fourth anthracycline dose revealed that HIF-1α protein was highly expressed in tumors from mice treated with vehicle or anthracycline (Figure 3C). By contrast, expression of mRNAs encoded by HIF-1 target genes was inhibited after anthracycline treatment (Figure 3D). Based on these results, we conclude that treatment of tumor-bearing mice with DNR or DXR inhibits HIF-1 transcriptional activity in tumor xenografts without affecting HIF-1α protein levels, as was observed in vitro.

### Anthracyclines Block PC-3 Prostate Cancer Xenograft Growth and Vascularization

As previously observed in HEK-293 (Figure S2A) and Hep3B-c1 (Figure 3C) cells, anthracyclines did not affect HIF-1α protein levels as determined by IB assay (Figure S5A) but blocked hypoxia-induced expression of HIF-1 reporter (Figure S5B) and target genes (Figure S5C) in PC-3 cells. Five million PC-3 cells were implanted subcutaneously into immunodeficient mice and when the tumors reached 100 mm^3^, the mice were randomized to receive either vehicle, DNR (0.5 or 1.5 mg/kg/day), or DXR (0.5 or1.5 mg/kg/day) by tail vein injection. Anthracycline administration blocked PC-3 xenograft growth as compared to the rapid tumor growth observed in vehicle-treated mice (Figure 4A).

**Figure 4.**
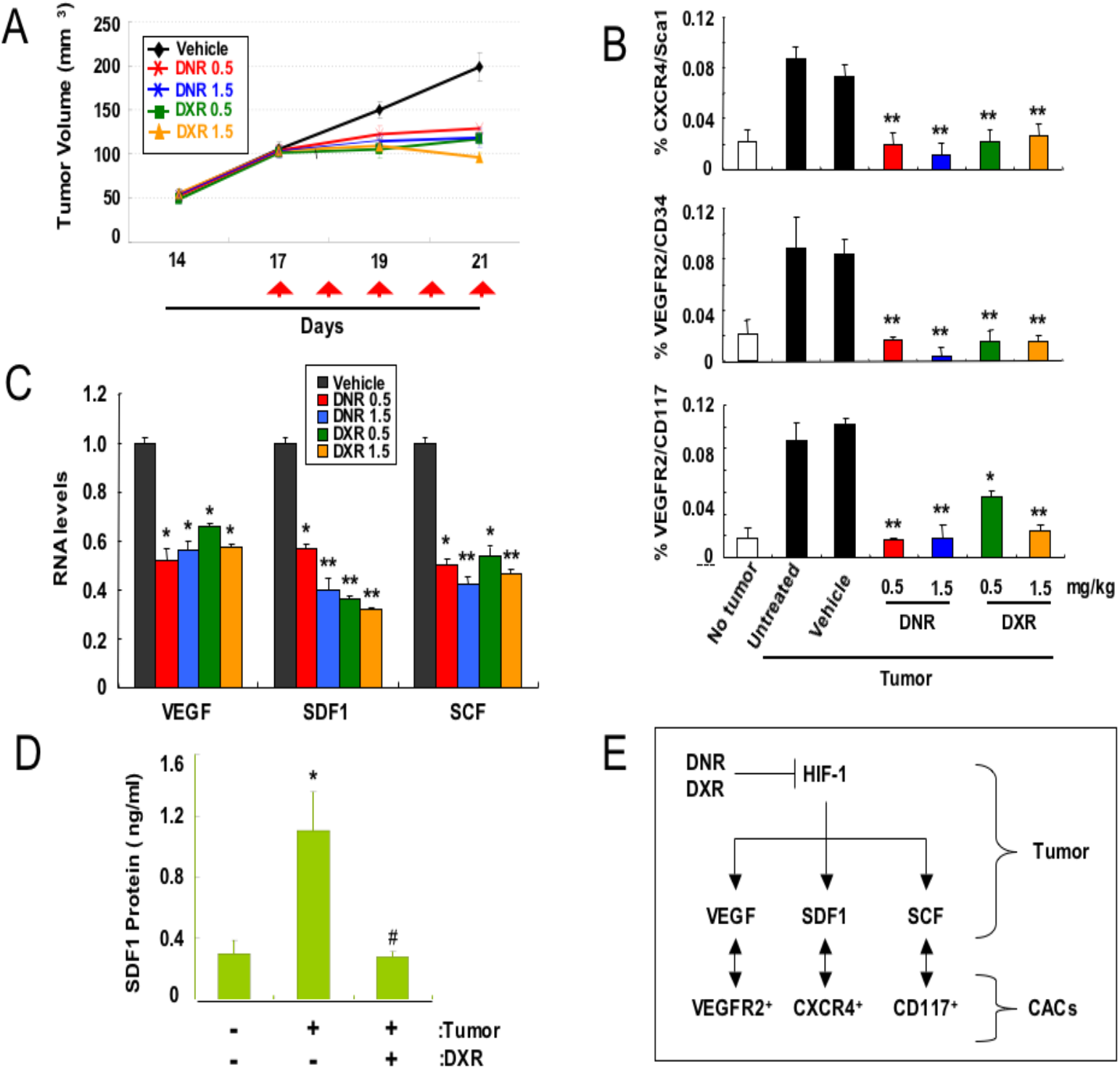
Anthracycline administration inhibits prostate tumor growth and CAC mobilization. (**A-C**) PC-3 prostate cancer xenografts were allowed to grow to 100 mm^3^ and mice were treated (red arrows) with vehicle (black), DNR (0.5 mg/kg/day, red; 1.5 mg/kg/day, blue), or DXR (0.5 mg/kg/day, green; 1.5 mg/kg/day, orange). Tumor volume was calculated on days 14, 17, 19 and 21 (**A**). Blood was collected four hours after treatment on day 21 and the percentage of CXCR4^+^/Sca1^+^, VEGFR2^+^/CD34^+^, and VEGFR2^+^/CD117^+^ circulating angiogenic cells (CACs) was determined by flow cytometry (mean ± SEM shown); *, *P* < 0.05; **, *P* < 0.01 (**B**). Tumors were harvested on day 21 and total RNA was isolated and analyzed by RT-qPCR using primers specific for VEGFA, SDF1, and SCF mRNA (mean ± SEM are shown; n = 4); *, *P* < 0.05; **, *P* < 0.01 (**C**). (**D**) Non-tumor bearing mice (-) and mice bearing prostate cancer xenografts of 100 mm^3^ (+) were treated with vehicle (-) or 0.5 mg/kg/day DXR (+) for 5 days. Blood samples were drawn four hours after the fifth dose and analyzed for SDF-1 α by ELISA (mean ± SEM shown; n =4. *, *P* < 0.01 vs. non-tumor-bearing mice; #, *P* < 0.01 vs. vehicle-treated tumor-bearing mice. (**E**) Administration of anthracycline to tumor-bearing mice inhibits HIF-1-dependent expression of angiogenic cytokines and CAC mobilization.

Tumors cannot grow beyond microscopic size without the establishment of a vasculature (27) and HIF-1 plays a critical role in the vascularization of tumor and normal tissue (1, 3-10). HIF-1 regulates the expression of multiple angiogenic growth factors, including VEGFA, SCF, and SDF-1 (4, 5, 28). VEGFA, SCF, and SDF-1 production by hypoxic cancer cells induces mobilization into the bloodstream of circulating angiogenic cells (CACs), which are a heterogeneous population of pro-angiogenic cells that home to tumors and mediate vascularization. By stimulating the expression of angiogenic cytokines, HIF-1 is essential for the homing of CACs (5, 6, 28, 29).

To investigate the effect of anthracyclines on CAC mobilization, tumor-bearing mice were treated daily with vehicle or anthracycline (Figure 4A) and 4 hours after the fifth dose, blood samples were collected for analysis of CACs by flow cytometry, based on co-expression of a progenitor cell marker [CD34, CD117 (also known as c-Kit), or Sca1 (stem cell antigen 1)] and either VEGFR2 or CXCR4, which are the receptors for VEGFA and SDF-1, respectively. CACs in peripheral blood were increased more than 4-fold in mice tumor-bearing as compared to non-tumor-bearing mice (Figure 4B). By contrast, DNR or DXR administration decreased CAC levels in the blood of tumor-bearing mice to levels that were similar to those in non-tumor-bearing mice.

To explore the molecular mechanism by which treatment with DNR or DXR blocked the mobilization of CACs into peripheral blood, we quantified angiogenic factor mRNA levels in tumor tissue. Anthracycline administration significantly decreased VEGFA, SDF-1, and SCF mRNA expression in PC-3 tumors (Figure 4C). We also measured circulating SDF-1 levels, which were increased in blood samples from vehicle-treated tumor-bearing mice; by contrast, SDF-1 levels in DXR-treated tumor-bearing mice were not significantly different from those measured in non-tumor-bearing mice (Figure 4D). Thus, DNR or DXR treatment blocks CAC mobilization in tumor-bearing mice by inhibiting the HIF-dependent expression of angiogenic factors (Figure 4E). Decreased expression of angiogenic cytokines and decreased CACs in peripheral blood were associated with a significant inhibition of angiogenesis in tumors from anthracycline-treated mice as determined by immunohistochemistry using antibodies against CD31, which is expressed by vascular endothelial cells, or a-smooth muscle actin (SMA), which is expressed by vascular pericytes and vascular smooth muscle cells (Figure 5).

**Figure 5.**
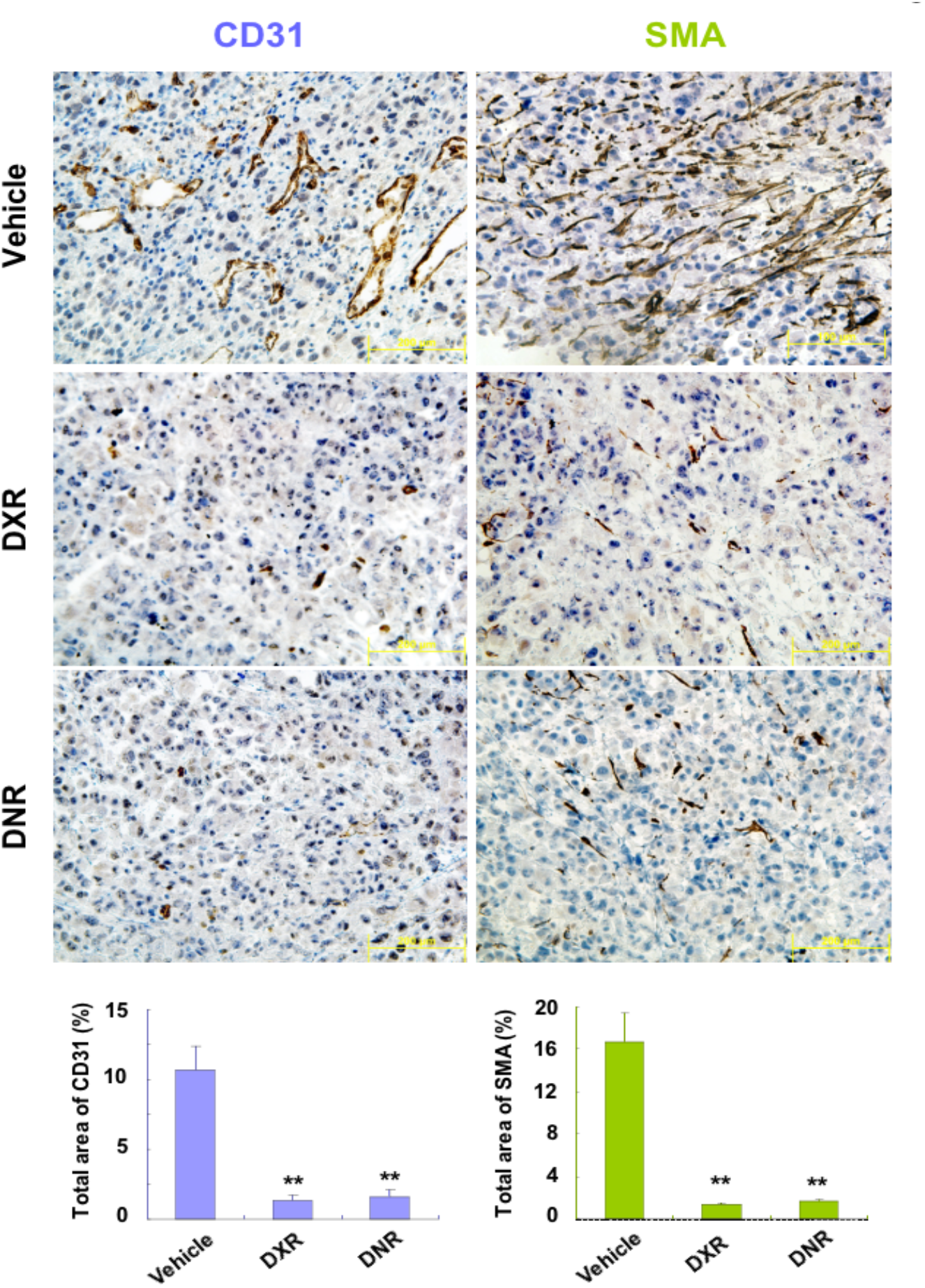
Anthracycline administration inhibits tumor angiogenesis. Prostate cancer xenografts were grown to a volume of 100 mm^3^ and mice were administered vehicle, DNR, or DXR (0.5 mg/kg/day) for 5 days. Immunohistochemistry was performed using antibody against CD31 or a-smooth muscle actin (SMA). The stained area (brown) in twenty fields in each of four tumors in each treatment group was quantified using ImageJ software (mean ± SEM shown). **, *P* < 0.01 versus vehicle.

## Discussion

In this study, we have shown that all four anthracycline chemotherapy agents tested inhibit HIF-1 activity, indicating a class effect. The therapeutic benefit of anthracyclines when administered at maximum tolerated dose is due to their ability to intercalate DNA and induce topoisomerase II-mediated strand breaks (26), but dose limiting toxicity in the bone marrow and gastrointestinal tract limits the efficacy of these drugs. Adriamycin (DXR) was reported to block tumor growth through anti-angiogenic effects when administered in small daily doses, which is known as metronomic therapy (26, 30, 31). The anthracycline aclacinomycin B was shown to inhibit expression of a HIF-1 reporter in Chinese hamster ovary cells; however, neither DXR nor DNR was found to inhibit HIF activity in that study (32). DXR treatment inhibited VEGFA mRNA expression in hypoxic ovarian cancer cells, but a mechanism of action was not determined (33). Anthracyclines have been shown to block the activity of several other transcription factors, including GATA4, AP1, and SP1 (34-36).

Our findings indicate that anthracyclines inhibit hypoxia-induced HIF-1 and HIF-2 binding to DNA in human cancer cells, as shown by ChIP assays; in contrast, anthracyclines do not affect HIF-1α protein expression, subunit heterodimerization, or TAD function. Anthracycline treatment of mice significantly inhibited PC-3 prostate cancer xenograft growth. These results complement immunohistochemistry data correlating increased HIF-1α levels in human prostate cancer biopsies with disease progression (37, 38).

Administration of anthracycline chemotherapy to tumor-bearing mice inhibited intratumoral expression of SCF, SDF-1 and VEGFA, which promote tumor angiogenesis by mobilizing CACs that express their cognate cell-surface receptors (CD117, CXCR4 and VEGFR2). Thus, inhibition of HIF-1 activity by anthracycline treatment leads to decreased SCF, SDF-1, and VEGFA expression, leading to impaired CAC mobilization, which results in decreased tumor vascularization and decreased tumor growth. It should be noted that while the present study was focused on the anti-angiogenic effects of HIF-1 inhibition, many other aspects of cancer biology are likely to be impacted, including cancer stem cell specification, immune evasion, metabolic reprogramming, and metastasis (3, 18, 19) and thereby contribute to the therapeutic effect of HIF-1 inhibition.

Our results have implications for the use of anthracyclines as anti-angiogenic agents in oncology. A general principle of targeted cancer therapy is that the target is selected based on evidence that increased expression or activity of the target is associated with disease progression. Thus, cancers in which HIF-1α overexpression in the diagnostic biopsy is associated with increased mortality, which includes prostate cancer (37, 38) and several other cancer types (18), are good candidates for inclusion in clinical trials of metronomic DXR or DNR therapy.

## Materials and Methods

### Cell Culture

HEK-293 and Hep3B-c1 cells were cultured in Dulbecco’s Modified Eagle Medium (DMEM; Mediatech). PC-3 cells were maintained in Roswell Park Memorial Institute-1640 medium (RPMI-1640; Invitrogen). Media were supplemented with 10% fetal calf serum (Hyclone), 100 U/ml penicillin, and 100 g/ml streptomycin (Invitrogen). Cells were incubated at 37 °C in 95% air and 5% CO_2_. For hypoxia treatment, cells were placed in a closed chamber flushed with a gas mixture containing 1% O_2_, 5% CO_2_, and 94% N2. DNR, DXR, EPI, and IDA (Sigma–Aldrich) were dissolved in DMSO at 1000x final concentration for use in cell-based studies.

### Drug Screen

Hep3B-c1 cells (10) were treated with the Hopkins Drug Library (20) or vehicle (DMSO), exposed to 20% or 1% O_2_ for 24 hours, lysed, and the FLuc:RLuc ratio was determined using the Dual Luciferase Assay system (Promega).

### Reverse Transcription and Quantitative Real-Time PCR (RT-qPCR) Assay

Total RNA was extracted using TRIzol (Invitrogen), treated with DNase (Ambion), a 1-μg aliquot was reverse-transcribed using the iScript cDNA synthesis system, and RT-qPCR was performed using iQ SYBR Green Supermix and the iCycler Real-time PCR detection system (Bio-Rad). Primer nucleotide sequences (Table S1) were determined using Beacon Designer software (Bio-Rad) and their specificity was analyzed by BLAST and PCR dissociation curve analysis. The expression level of each mRNA was normalized to the expression of 18S rRNA in the same sample.

### IB Assays

Whole cell lysates were prepared in lysis buffer [50 mM Tris Cl (pH 7.5), 150 mM NaCl, 0.1% Nonidet P-40, 1 mM DTT, 1 mM Na3VO4, 10mM NaF, and protease inhibitor mixture] and IB assays were performed as previously described (39) using antibody against either FLAG (Sigma–Aldrich), HIF-1α, or β-actin (Santa Cruz Biotechnology).

### GST Pulldown Assay

GST-HIF-1β (amino acid residues 11–510) was generated by PCR amplification from cDNA (primers shown in Table S2). The PCR product was inserted into pGEX- 5X-1 (GE Healthcare). GST fusion proteins were purified as described (39). Whole cell lysates were prepared from HEK-293 cells transfected with an expression vector encoding FLAG-HIF-1α using lysis buffer. An 800-μg aliquot of lysate and 4 μg of the purified GST-HIF-1β fusion protein or GST alone were incubated in the presence of 5 μM DNR, DXR, EPI, or IDA, or vehicle control (DMSO), overnight at 4 °C. A 30-μl aliquot of glutathione–Sepharose 4B beads (GE Healthcare) was added and incubated for 3 hours at 4 °C, followed by washing with lysis buffer, and elution of bound proteins using Laemmli buffer (BioRad).

### GAL4 Reporter Assay

HEK-293 cells were seeded at fifty thousand cells per well in a 24-well plate and incubated for 20 hours in a standard tissue culture incubator. Cells were co-transfected with 150 ng of pGAL4-E1b-Luc and 200 ng of expression vector encoding the GAL4 DBD alone or fused to HIF-1α coding sequences (21) using FuGENE-6. pSV-RLuc was used as an internal control. Cells were exposed to 20% or 1% O_2_ for 24 hours in the presence of vehicle or 5 μM DNR, DXR, EPI, or IDA, then lysed and the FLuc:RLuc ratio was determined.

### ChIP Assay

HEK-293 cells were treated with 1 μM DNR, DXR, EPI, or IDA and exposed to 20% or 1% O_2_ for 24 hours. ChIP was performed with the ChIP Assay kit (Upstate–Cell Signaling Solution) using rabbit polyclonal anti-HIF-1a or anti-HIF-2α antibodies (Novus Biologicals) and normal rabbit IgG as a negative control (Santa Cruz Biotechnology). *VEGFA* and *PDK1* gene sequences were detected by PCR using the oligonucleotide primers listed in Table S3.

### Xenograft Assays

Male 5-week-old severe combined immunodeficiency mice were purchased from the National Cancer Institute. DXR, DNR, and saline used for animal studies were purchased from the research pharmacy of the Johns Hopkins Hospital. Mice were implanted subcutaneously with 5 × 10^6^ Hep3B-c1 or PC-3 cells suspended in 0.2 ml of DMEM or RPMI-1640, respectively. The mice were examined every 2-3 days for measurement of body weight and tumor volume (V), which was calculated according to the following formula: V = length × width × height × 0.52.

### Bioluminescent Imaging

Mice bearing Hep3B-c1 xenografts received an intraperitoneal injection of 3 mg of D-luciferin firefly (Caliper Life Science) dissolved in 0.2 ml of PBS. The mice were analyzed using the Xenogen IVIS 200 optical imaging device in the Johns Hopkins Small Animal Molecular Imaging Center.

### Flow Cytometry

A 100-μl aliquot of anti-coagulated peripheral blood was incubated with 1 μg each of fluorescein isothiocyanate- and phycoerythrin-conjugated monoclonal antibodies against CXCR4 and Sca1, VEGFR2 and CD34, or VEGFR2 and CD117, respectively (BD Biosciences). Erythrocytes were lysed by addition of 0.8% NH4Cl and 10 mM EDTA (Stem Cell Technologies). Cells were collected by centrifugation for 5 min at 400 × *g* and analyzed using a FACScan flow cytometer (BD Biosciences), which was equipped with an argon laser emitting at 488 nm. From 100,000 events, the percentage of cells showing dual fluorescence was determined.

### Tumor Immunohistochemistry

PC-3 tumors were fixed in 10% formalin, embedded in paraffin, and 5-μm sections were prepared, mounted on positively charged slides, hydrated through xylene and graded ethanol, and washed with PBS. Endogenous peroxidase activity was quenched using 3% hydrogen peroxide in methanol and nonspecific binding was blocked by incubation with blocking solution (BioGenex). CD31 (1:50; BD Pharmingen) or SMA (1:50; Sigma) antibodies were applied for 1 hour, the slides were washed with PBS containing 0.1% Tween 20, incubated with biotinylated secondary antibody (1:1,000; Vector Laboratories) for 1 hour, antibody staining was visualized with Vectastain Elite ABC immunoperoxidase system (Vector Laboratories), and sections were counterstained with hematoxylin. The CD31- or SMA-positive area was quantified (40 × objective, 20 fields from each slide) using ImageJ software (National Institutes of Health).

### SDF-1 ELISA

Blood was collected from mice, allowed to clot, and centrifuged at 2,000 × *g* for 20 min. SDF-1α levels were determined in duplicate for each sample by ELISA (R&D Systems). Optical density was measured at 450 nm and 570 nm (correction) using an AD340 plate reader (Beckman Coulter). SDF-1α levels were calculated by linear regression from a standard curve.

## ACKNOWLEDGMENTS

We thank Drs. KangAe Lee and Hong Wei for their contributions to this work. We are grateful to Karen Padgett (Novus Biologicals) for providing anti-FLAG and anti-HIF-2α antibodies. This work was supported by the Armstrong Family Foundation, Flight Attendant Medical Research Institute, and the Johns Hopkins Institute for Cell Engineering.

## Supplemental Figures and Tables

**Figure S1.**
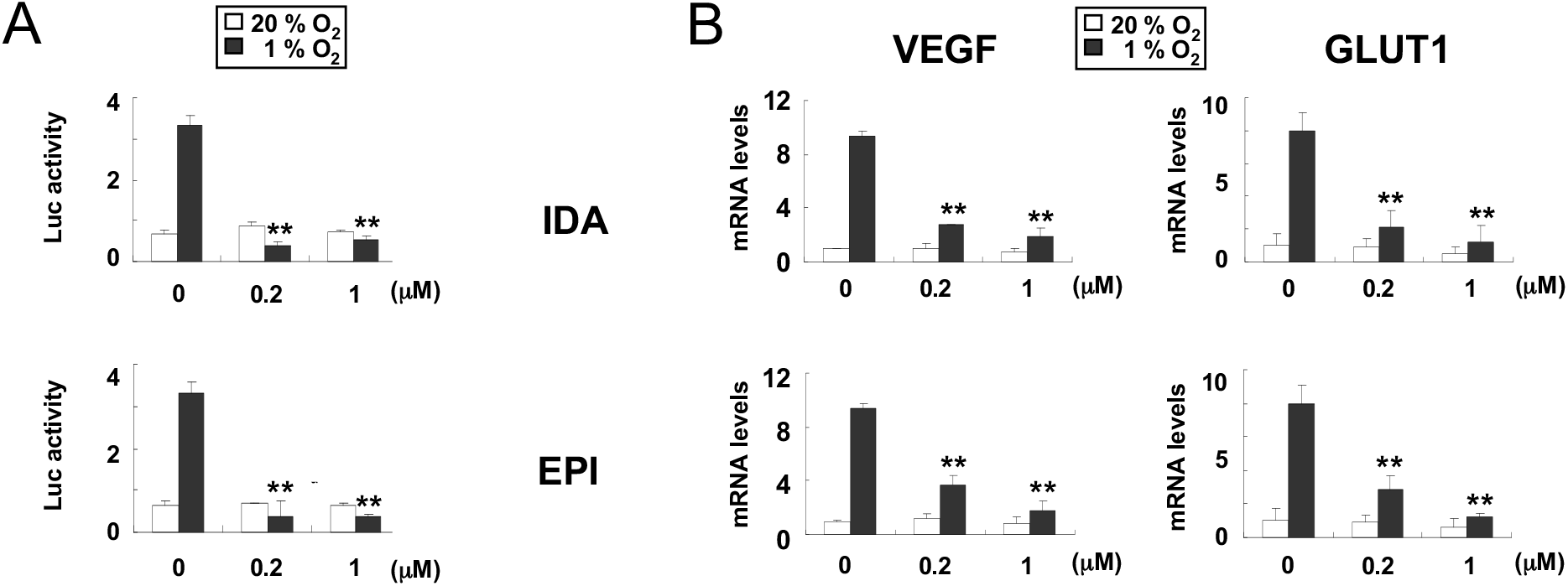
Idarubicin (IDA) and epirubicin (EPI) inhibit HIF-1 activity. (**A**) Hep3B-c1 cells were treated with the indicated concentration of IDA *(top panel)* or EPI *(bottom panel)* and incubated at 20% O_2_ (white bars) or 1% O_2_ (black bars) for 24 hours. Hypoxia-induced luciferase activity (FLuc:RLuc ratio) was measured in cell lysates. (**B**) HEK-293 cells were exposed to 20% O_2_ (white bars) or 1% O_2_ (black bars) for 24 hours in the presence of the indicated concentration of IDA (*top panels)* or EPI *(bottom panels).* Total RNA was isolated and levels of VEGFA mRNA *(left panels)* and GLUT1 mRNA *(right panels)* were determined using RT-qPCR. *, *P* < 0.05; **, *P* < 0.01.

**Figure S2.**
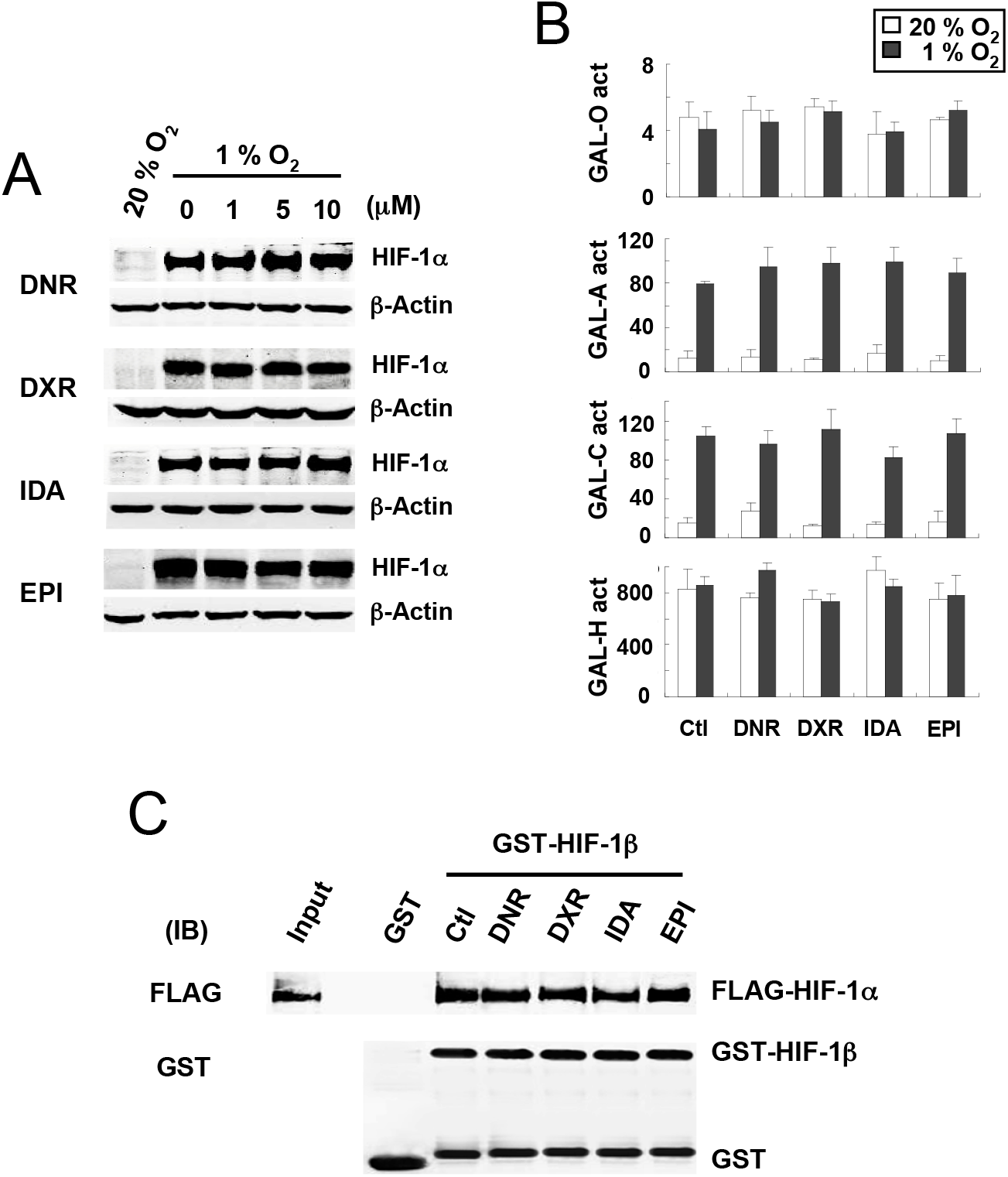
Effect of anthracyclines on HIF-1α protein expression, HIF-1α transactivation domain (TAD) function and HIF-1 DNA binding activity. (**A**) HIF-1α and β-Actin protein were analyzed by IB assays of lysates of HEK-293 cells that were cultured at 1% O_2_ for 20 hours in the presence of the indicated concentration of the indicated anthracycline. (**B**) HIF-1α TAD function was analyzed using GAL4/HIF-1α fusion proteins to drive FLuc expression in HEK-293 cells that were treated with vehicle (Con) or the indicated anthracycline at a concentration of 5 μM and exposed to 1% O_2_ (black bars) or 20% O_2_ (white bars) for 24 hours. The data are presented as mean ± SEM ratio of FLuc:RLuc activity from 3 independent transfection experiments. (**C**) Heterodimerization of HIF-1α and HIF-1β subunits was determined by incubating a GST-HIF-1β fusion protein with lysate from HEK-293 cells expressing FLAG-tagged HIF-1α in the presence of 5 μM of the indicated anthracycline. Proteins were pulled down with glutathione-Sepharose-4B beads and IB assays were performed with the indicated antibodies.

**Figure S3.**
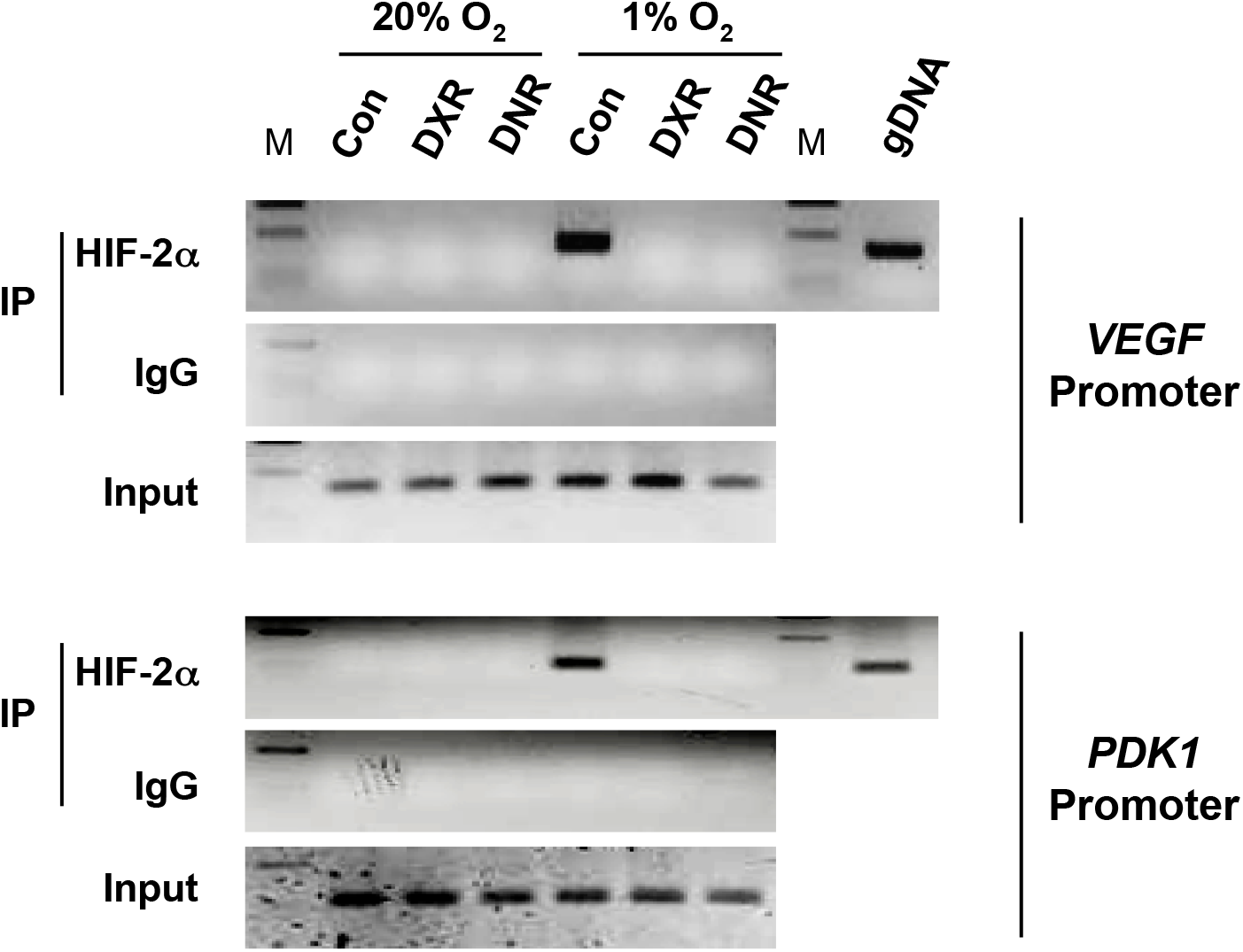
Anthracyclines inhibit HIF-2 DNA-binding activity as determined by chromatin immunoprecipitation assay. HEK-293 cells were exposed to the indicated O_2_ concentration for 20 hours in the presence of vehicle (Con) or the indicated anthracycline (1 μM). Aliquots of chromatin were incubated with HIF-2α antibody or rabbit IgG. Immunoprecipitated DNA was used as template to amplify *VEGF* and *PDK1* promoter sequences that encompass a HIF binding site. PCR products were analyzed by 2% agarose gel electrophoresis and ethidium bromide staining. An aliquot of chromatin was reserved before immunoprecipitation for PCR (Input). Purified genomic DNA was also subject to PCR (gDNA). M, DNA ladder.

**Figure S4.**
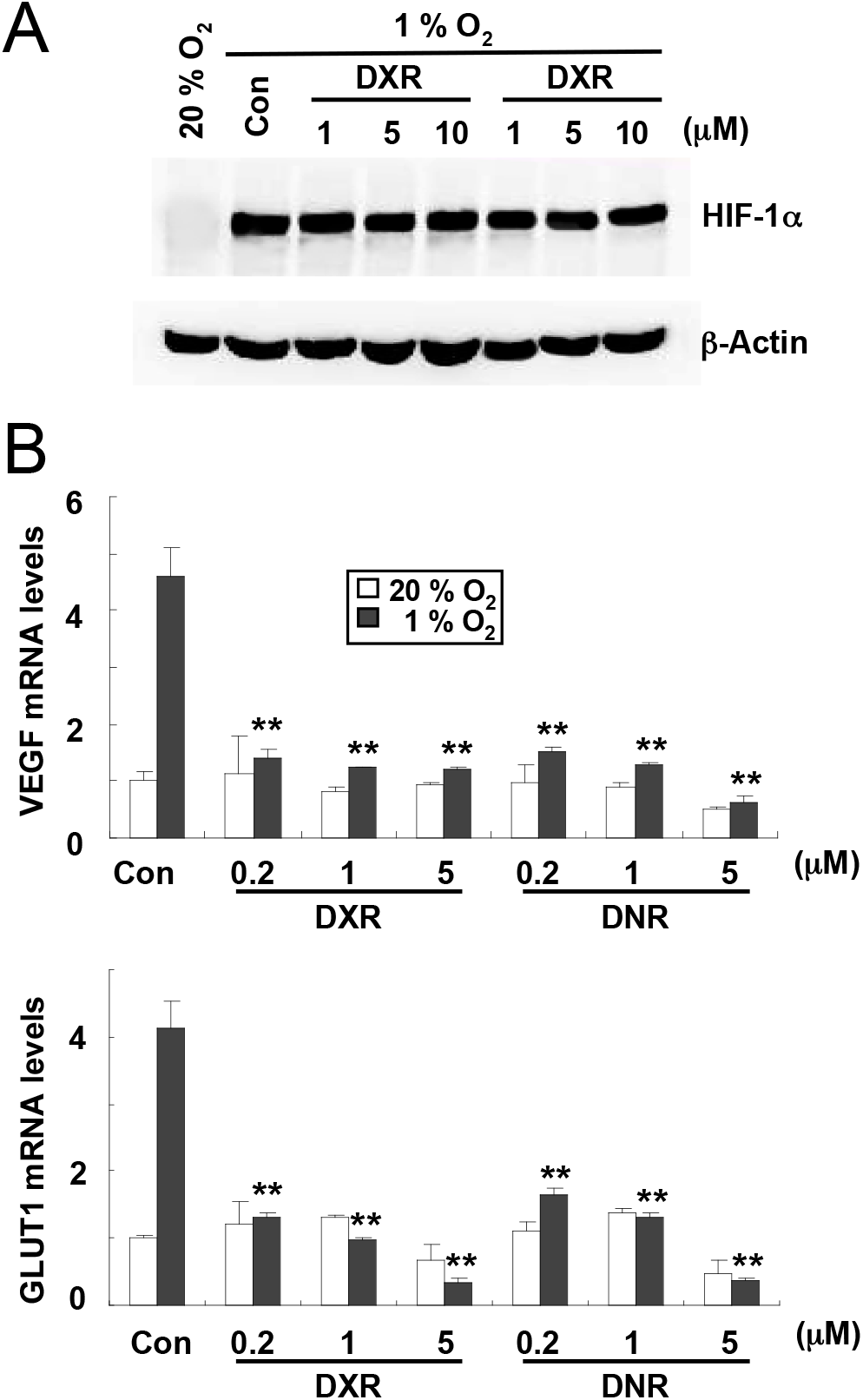
Effect of anthracycline treatment on HIF-1α expression and HIF-1 target gene expression. (**A**) HIF-1α and β-actin expression was analyzed by IB assays of lysates prepared from hypoxic Hep3B-c1 cells that were exposed to 1% O_2_ for twenty hours and treated with vehicle control (Con) or the indicated concentration of anthracycline. (**B**) mRNA expression was analyzed in Hep3B- c1 cells cultured in 1% O_2_ (black bars) or 20% O_2_ (white bars) in the presence of vehicle (Con) or the indicated anthracycline. The data are presented as mean + SD (n = 4). *\*\*P* < 0.01 vs Con.

**Figure S5.**
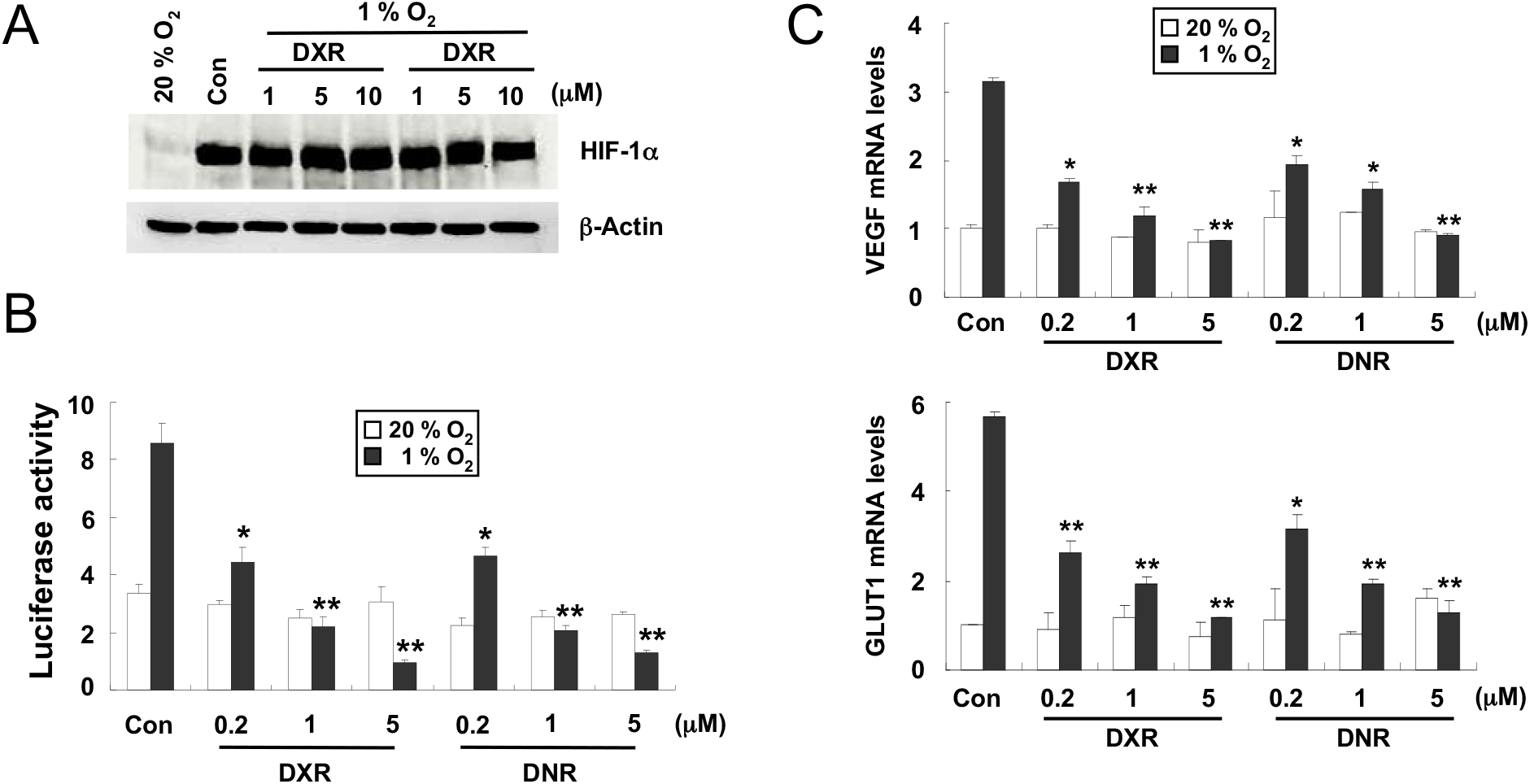
Analysis of anthracycline effects in prostate cancer cells. **(A)** Analysis of HIF-1α and β-actin expression. PC-3 cells were cultured in 1% O_2_ for twenty hours in the presence of vehicle (Con) or the indicated anthracycline concentration and IB assays were performed. (**B**) Analysis of HIF-1- dependent luciferase activity. PC-3 cells were co-transfected with HIF-1-dependent FLuc and control RLuc reporter plasmids and treated with anthracycline for 24 hours at 20% O_2_ (white bars) or 1% O_2_ (black bars). Cells were lysed and the FLuc:RLuc ratio was determined. (**C**) mRNA levels were measured in cells treated with anthracycline at 20% O_2_ (white bars) or 1% O_2_ (black bars) O_2_ for 24 hours. The data are presented as mean ± SEM (n = 4). * *P* < 0.05; **, *P* < 0.01.

**Table S1.**
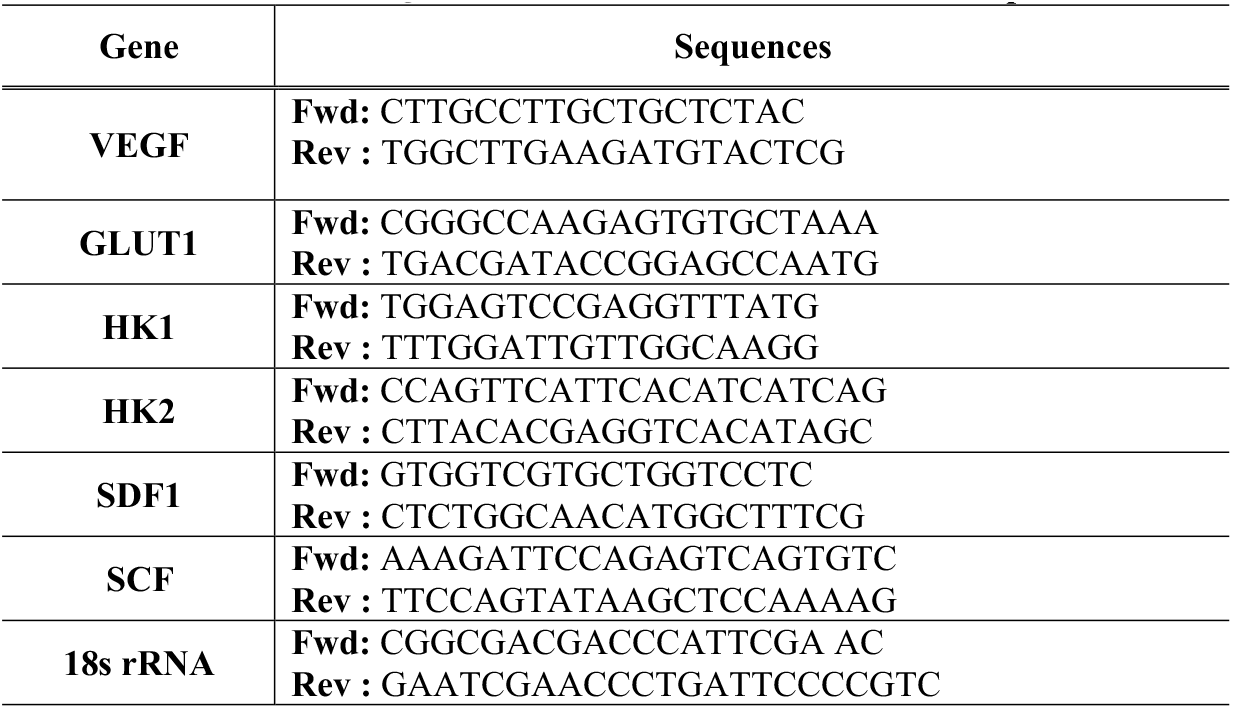
Primers used for Quantative Real-Time Reverse-Transcription PCR.

**Table S2.**
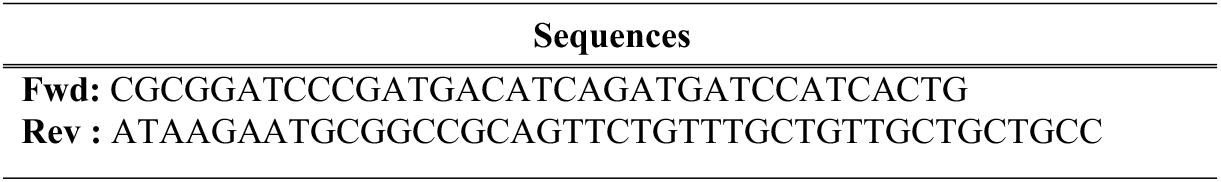
Primers Used for the Amplification of HIF-1β_11-510_.

**Table S3.**
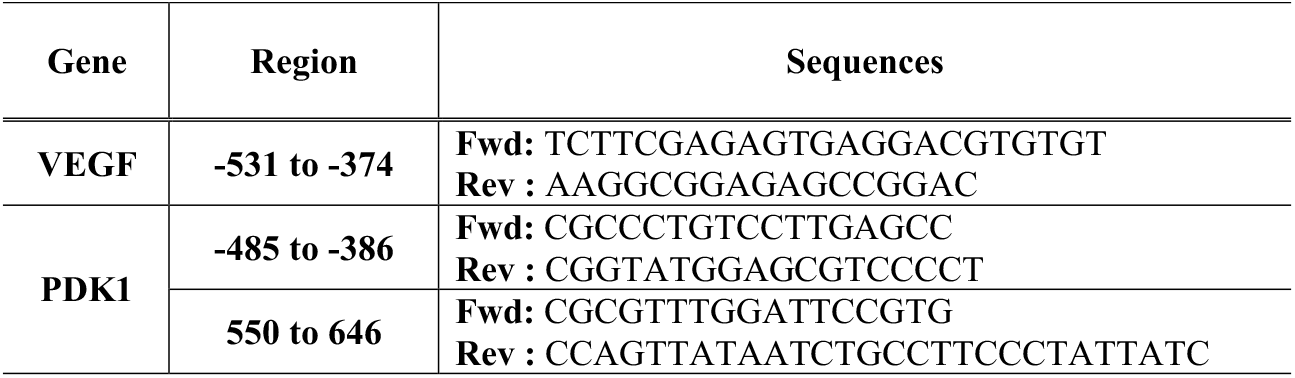
Primers Used for Chromatin Immunoprecipitation Assay.

## References

1. Brahimi-Horn MC, Chiche J, Pouyssegur J. Hypoxia and cancer. J Mol Med 2007;26:225–239.

2. Vaupel P, Höckel M, Mayer A. Detection and characterization of tumor hypoxia using pO_2_ histography. Antioxid Redox Signal 2007;9:1221–1235.

3. Semenza GL. The hypoxic tumor microenvironment: A driving force for breast cancer progression. Biochim Biophys Acta 2016;1863:382–391.

4. Forsythe JA, Jiang BH, Iyer NV, et al. Activation of vascular endothelial growth factor gene transcription by hypoxia-inducible factor 1. Mol Cell Biol 1996;16:4604–4613.

5. Bosch-Marce M, Okuyama H, Wesley JB, et al. Effects of aging and hypoxia-inducible factor 1 activity on angiogenic cell mobilization and recovery of perfusion after limb ischemia. Circ Res 2007;101:1310–1318.

6. Hirota K, Semenza GL. Regulation of angiogenesis by hypoxia-inducible factor 1. Crit Rev Oncol Hematol 2006;59:16–26.

7. Iyer NV, Kotch LE, Agani F, et al. Cellular and developmental control of O2 homeostasis by hypoxia-inducible factor 1α. Genes Dev 1998;12:149–162.

8. Lee K, Zhang H, Qian DZ, et al. Acriflavine inhibits HIF-1 dimerization, tumor growth, and vascularization. Proc Natl Acad Sci USA 2009;106:17910–17915.

9. Stoeltzing O, McCarty MF, Wey JS, et al. Role of hypoxia-inducible factor 1α in gastric cancer cell growth, angiogenesis and vessel maturation. J Natl Cancer Inst 2004;96:946–956.

10. Zhang H, Qian DZ, Tan YS, et al. Digoxin and other cardiac glycosides inhibit HIF-1α synthesis and block tumor growth. Proc Natl Acad Sci USA 2008;105:19579–19586.

11. Semenza GL. HIF-1 mediates metabolic responses to intratumoral hypoxia and oncogenic mutations. J Clin Invest 2013;123:3664–3671.

12. Semenza GL, Jiang BH, Leung SW, et al. Hypoxia response elements in the aldolase A, enolase 1, and lactate dehydrogenase A gene promoters contain essential binding sites for hypoxia-inducible factor 1. J Biol Chem 1996;271:32529–32537.

13. Wang GL, Semenza GL. Purification and characterization of hypoxia-inducible factor 1. J Biol Chem 1995;270:1230–1237.

14. Jiang BH, Semenza GL, Bauer C, et al. Hypoxia-inducible factor 1 levels vary exponentially over a physiologically relevant range of O_2_ tension. Am J Physiol 1996;271:C1172–C1180.

15. Jaakkola P, Mole DR, Tian YM, et al. Targeting of HIF-a to the von Hippel-Lindau ubiquitylation complex by O_2_-regulated prolyl hydroxylation. Science 2001;292:468–472.

16. Maxwell PH, Wiesener MS, Chang GW, et al. The tumor suppressor protein VHL targets hypoxia-inducible factors for oxygen-dependent proteolysis. Nature 1999;399:271–275.

17. Lando D, Peet DJ, Gorman JJ, et al. FIH-1 is an asparaginyl hydroxylase enzyme that regulates the transcriptional activity of hypoxia-inducible factor. Genes Dev 2002;16:1466–1471.

18. Wicks EE, Semenza GL. Hypoxia-inducible factors: cancer progression and clinical translation. J Clin Invest 2022;132:e159839.

19. Semenza GL. Pharmacologic targeting of hypoxia-inducible factors. Annu Rev Pharmacol Toxicol 2019;59:379–403.

20. Chong CR, Xu J, Lu J, et al. Inhibition of angiogenesis by the antifungal drug itraconazole. ACS Chem Biol 2007;2:263–270.

21. Jiang BH, Zheng JZ, Leung SW, et al. Transactivation and inhibitory domains of hypoxia-inducible factor 1α: modulation of transcriptional activity by oxygen tension. J Biol Chem 1997;272:19253–19260.

22. Mahon PC, Hirota K, Semenza GL. FIH-1: a novel protein that interacts with HIF-1α and VHL to mediate repression of HIF-1 transcriptional activity. Genes Dev 2001;15:2675–2686.

23. Chaires JB, Herrera JE, Waring MJ. Preferential binding of daunomycin to 5’ATCG and 5’ATGC sequences revealed by footprinting titration experiments. Biochemistry 1990;29:6145–6153.

24. Priebe W, Fokt I, Przewloka T, et al. Exploiting anthracycline scaffold for designing DNA-targeting agents. Methods Enzymol 2001;340:529–555.

25. Hurley LH. DNA and its associated processes as targets for cancer therapy. Nat Rev Cancer 2002;2:188–200.

26. Quesada AJ, Nelius T, Yap R, et al. In vivo up-regulation of CD95 and CD95L causes synergistic inhibition of angiogenesis by TSP1 peptide and metromomic doxorubicin treatment. Cell Death Differ 2005;12:649–658.

27. Folkman J. Tumor angiogenesis: therapeutic implications. N Engl J Med 1971;285:1182–1186.

28. Ceradini DJ, Kulkarni AR, Callaghan MJ, et al. Progenitor cell trafficking is regulated by hypoxic gradients through HIF-1 induction of SDF-1. Nat Med 2004;10:858–864.

29. Du R, Lu KV, Petritsch C, et al. HIF-1a induces the recruitment of bone marrow-derived vascular modulatory cells to regulate tumor angiogenesis and invasion. Cancer Cell 2008;13:206–220.

30. Drevs J, Fakler J, Eisele S, et al. Antiangiogenic potency of various chemotherapeutic drugs for metronomic chemotherapy. Anticancer Res 2004;24:1759–1763.

31. Kerbel RS, Kamen BA. The anti-angiogenic basis of metronomic chemotherapy. Nat Rev Cancer 2004;4:423–436.

32. Yamazaki Y, Hasebe Y, Egawa K, et al. Anthracyclines, small-molecule inhibitors of hypoxia-inducible factor-1a activation. Biol Pharm Bull 2006;29:1999–2003.

33. Duyndam MC, van Berkel MPA, Dorsman JC, et al. Cisplatin and doxorubicin repress vascular endothelial growth factor expression and differentially down-regulate hypoxia-inducible factor 1 activity in human ovarian cancer cells. Biochem Pharmacol 2007;74:191–201.

34. Kim Y, Ma AG, Kitta K, et al. Anthracycline-induced suppression of GATA-4 transcription factor: implication in the regulation of cardiac myocyte apoptosis. Mol Pharmacol 2003;63:368–377.

35. Szulawska A, Gniazdowski M, Czyz M. Sequence specificity of formaldehyde-mediated covalent binding of anthracycline derivatives to DNA. Biochem Pharmacol 2005;69:7–18.

36. Mansilla S, Portugal J. Sp1 transcription factor as a target for anthracyclines: effects on gene transcription. Biochimie 2008;90:976–987.

37. Vergis R, Corbishley CM, Norman AR, et al. Intrinsic markers of tumor hypoxia and angiogenesis in localized prostate cancer and outcome of radical treatment: a retrospective analysis of two randomized radiotherapy trials and one surgical cohort study. Lancet Oncol 2008;9:342–351.

38. Ranasinghe WKB, Xiao L, Kovac S, et al. The role of hypoxia-inducible factor 1α in determining the properties of castrate-resistant prostate cancers. PLoS One 2013;8:e54251.

39. Baek JH, Liu YV, McDonald KR, et al. Spermidine/spermine-N1-acetyltransferase 2 is an essential component of the ubiquitin ligase complex that regulates hypoxia-inducible factor 1α. J Biol Chem 2007;282:23572–23580.

